# Attention-based deep multiple instances learning for classifying circular RNA and other long non-coding RNA

**DOI:** 10.1101/2021.09.01.458499

**Authors:** Yunhe Liu, Qiqing Fu, Xueqing peng, Chaoyu Zhu, Gang Liu, Lei Liu

## Abstract

Circular RNA (circRNA) is a distinguishable circular formed long non-coding RNA (lncRNA), which has specific roles in transcriptional regulation, multiple biological processes. The identification of circRNA from other lncRNA is necessary for relevant research. In this study, we designed attention-based multi-instance learning (MIL) network architecture, which can be fed with raw sequence, to learn the sparse features in sequences and accomplish the identification task for circRNAs. The model outperformed previously reported models. Following the effectiveness validation of the attention score by the handwritten digit dataset, the key sequence loci underlying circRNAs recognition were obtained based on the corresponding attention score. Moreover, the motif enrichment analysis of the extracted key sequences identified some of the key motifs for circRNA formation. In conclusion, we designed a deep learning network architecture suitable for gene sequence learning with sparse features and implemented to the circRNA identification, and the network has a strong representation capability with its indication of some key loci.

## Introduction

Non-coding RNAs (ncRNAs), referring to RNAs without protein-coding potential, account for the majority of RNAs. It’s generally recognized that lncRNA (long non-coding RNA) is a kind of ncRNAs that is longer than 200 nucleotides, which is distinguished from other smaller ncRNA species such as miRNAs and siRNAs. lncRNA has complex biological functions such as transcriptional regulation and post-transcriptional control^[1–3]^. Circular RNA (circRNA) is a closed lncRNA formed by covalently closed loops. Based on current researches, circRNAs are more stable than mRNAs and play a major role as a microRNA activity modulator. CircRNAs are also correlated with the development of multiple diseases^[3–5]^, and can be used for disease biomarkers^[6, 7]^. Therefore, it is vital to detect circular RNAs.

Currently, some computational approaches to distinguish circRNA from lncRNA^[8–10]^ have been developed with different frameworks. For example, CirRNAPL^[11]^ adopted the extreme learning machine based on the particle swarm optimization algorithm. CircLGB utilized a LightGBM classifier^[12]^. Based on an End-to-End deep learning framework, circDeep^[13]^ fused an RCM, ACNN-BLSTM sequence, and a conservation descriptor into high-level abstraction descriptors, and achieved an improvement with higher accuracy compared with exiting tools.

For these models mentioned above, the input of the model was not the raw sequence, but often the relevant features extracted from the predicted secondary structure^[9, 13]^. For circDeep, a deep learning framework utilized complementary structure^[14]^ and conservation features. Its sequence input part did not use the full sequence and underwent a triplet transformation^[15]^, either. It is important to find a deep learning framework that is suitable for sequence input as well as sequence learning, to facilitate the utilization of algorithms and take advantage of the information in sequence.

The characteristics of RNA sequences have quite different from the other sequence data like word language. We organized the differences into 3 main points. First, the RNA sequences are a combination of multiple meaningful and meaningless sequences, where the meaningful units are embedded into the entire background sequences, not like the words that form a certain grammatical structure in order^[16]^. An RNA typically has a large variety of functions enabled by meaningful units, such as the ability to form high-level structures and to recruit other components^[17]^. While learning models tend to have singular learning objectives, such as distinguishing circular RNA, which results in the meaningful units for learning model is sparse in the long sequence. Second, the length of different RNA varies greatly^[18]^, spans from 100 to 1,000,000 nt, suggesting that the density of the meaningful units also varies considerably. Third, the character component of the RNA sequence is relatively simple, which only contains four characters (ATGC) and the single character is meaningless. On the other hand, the composition and length of meaningful components are unknown, so the input data for learning is character instead of a meaningful word.

To address the problem mentioned above, we designed an attention-based deep encoder MIL (multiple instance learning) model (Circ-ATTEN-MIL). The MIL structure is suitable for learning sparse features^[19, 20]^, and the attention-based pooling layer can discover similarities between instances and has a stronger representation capability^[21]^. We applied this deep network structure to learn the identification task and achieved better accuracy and extracted high attention sequences to enrich motif, which shed light on studies regarding RNA circligase.

## Method

### Data source

We extracted circRNAs sequences from the circRNADb database^[22]^ and other lncRNAs sequences from the GENCODE database^[23]^ (lincRNA, antisense, processed transcript, sense intronic, and sense overlapping) respectively. After removed sequences shorter than 200 nucleotides, we got 31939 circRNAs and 19722 other lncRNAs. The circRNA sequences were regarded as positive samples. We randomly divided the dataset into a training set (75%), validation set (10%), and test set (15%).

### Instances extraction by sliding window

An RNA sequence was regarded as a bag, and instances were extracted from the sequence. For each full-length sequence, we connected the head (5’ end) and tail (3’ end) of the sequence, set the slider window size and the sliding step, and made the slider move from the head. For each step, the sequence contained in the window was extracted as an instance, until the slider moved out of the tail of the sequence (Illustrated in Fig.1). For a sequence of a certain length, the number of instances can be calculated by the following formula.

**Figure 1.**
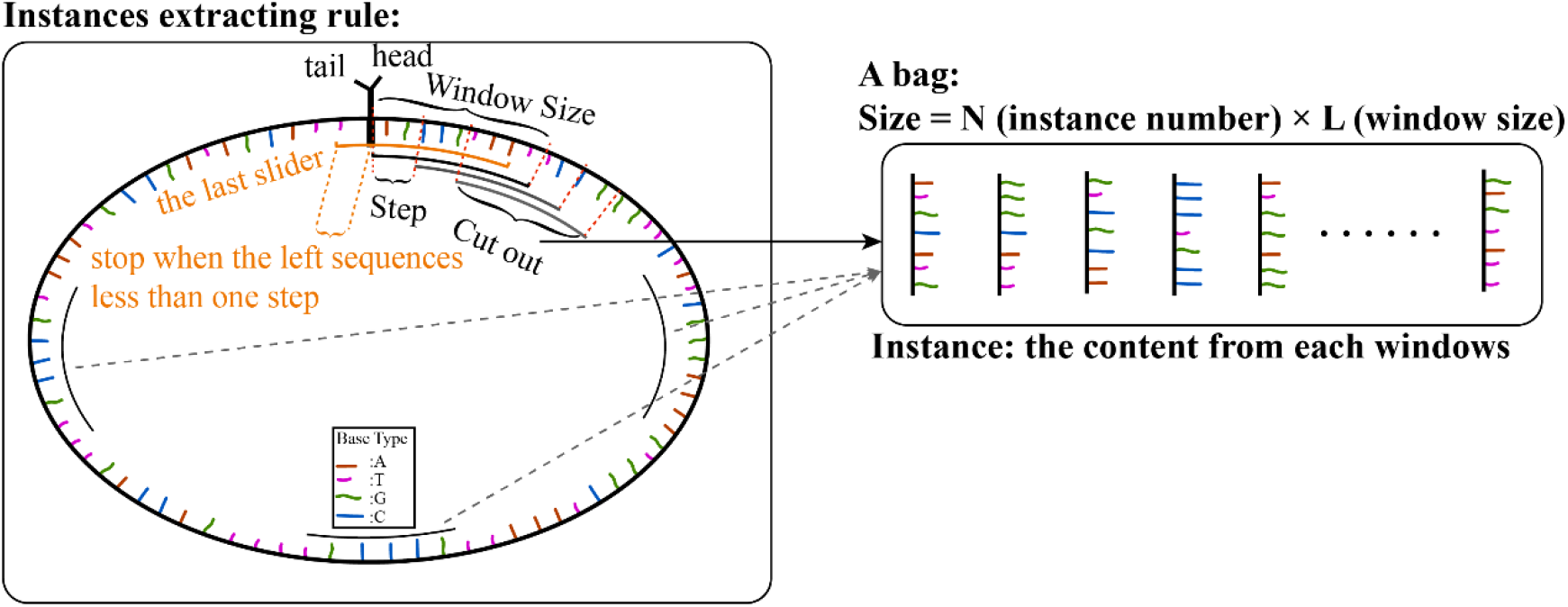
Illustrations of instance extraction from full RNA sequence.

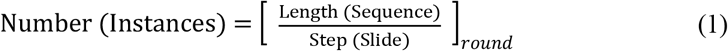

### Model structure

The network structure was represented in Figure 2. We employed the encoder structure of the seq2seq model^[24]^ here as the instance feature extractor. The embedding layer^[25]^ was employed to represent bases (15 (A, T, G, C, N, H, B, D, V, R, M, S, W, Y, K) → 4 (representative dimension))_∘_ The encoder used a bi-directional RNN structure, which given equal attention to the head and the tail of the instance, and the output was a context vector^[26]^ to represent the feature of the instance. And subsequently, through the MIL layer, the features of all instances were scored and aggregated jointly to determine the type of the bag^[20, 21, 27]^.

**Figure 2.**
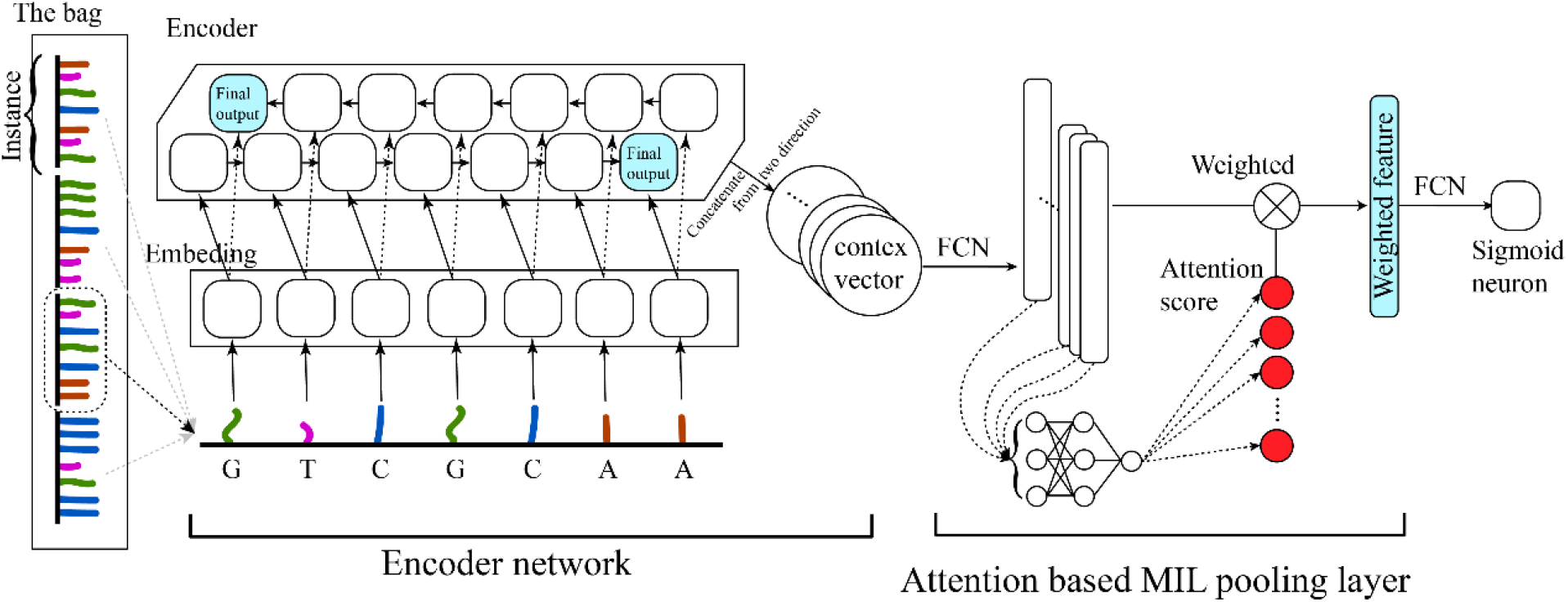
Illustrations of attention-based deep encoder MIL model structure (Circ-ATTEN-MIL).

### Attention mechanism as the MIL pooling

Reference to previous work of pooling layer structure, we selected the attention-based pooling structure, which exhibited better aggregation and representation capacity^[21]^. It was assumed that the feature extracted by encoder were C = {**c**_1_, …, **c**_*k*_}, and its corresponding attention weights were ***α*** = {*α*_1_, …, *α*_*k*_}, which could be formulated as follow.

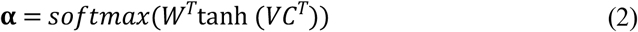

Where W _∈_ R^*L*×1^ and V _∈_ R^*L*×*M*^. The attention-base structure allowed to discover the similarity between different instances and made the network have better representability. After the encoder feature was weighted by the attention scores, the probability of determination was output via a sigmoid neuron through a fully connected layer.

### Handling of handwritten numbers dataset

The handwritten numbers dataset was used to verify the representational power of the attention score. Each number figure (size = 28×28) was served as an instance. A bag contained more than 16 instances. For each instance, we treated the image as a sequence containing 28 characters, and each with a representation dimension of 28, for feeding into the network (Circ-ATTEN-MIL; the embedding layer in encoder block was removed in this task) (Fig.3). A bag is positive when it contains the determining number (Two modes were set: determining number is 0; determining number are 0, 1, 3).

**Figure 3.**
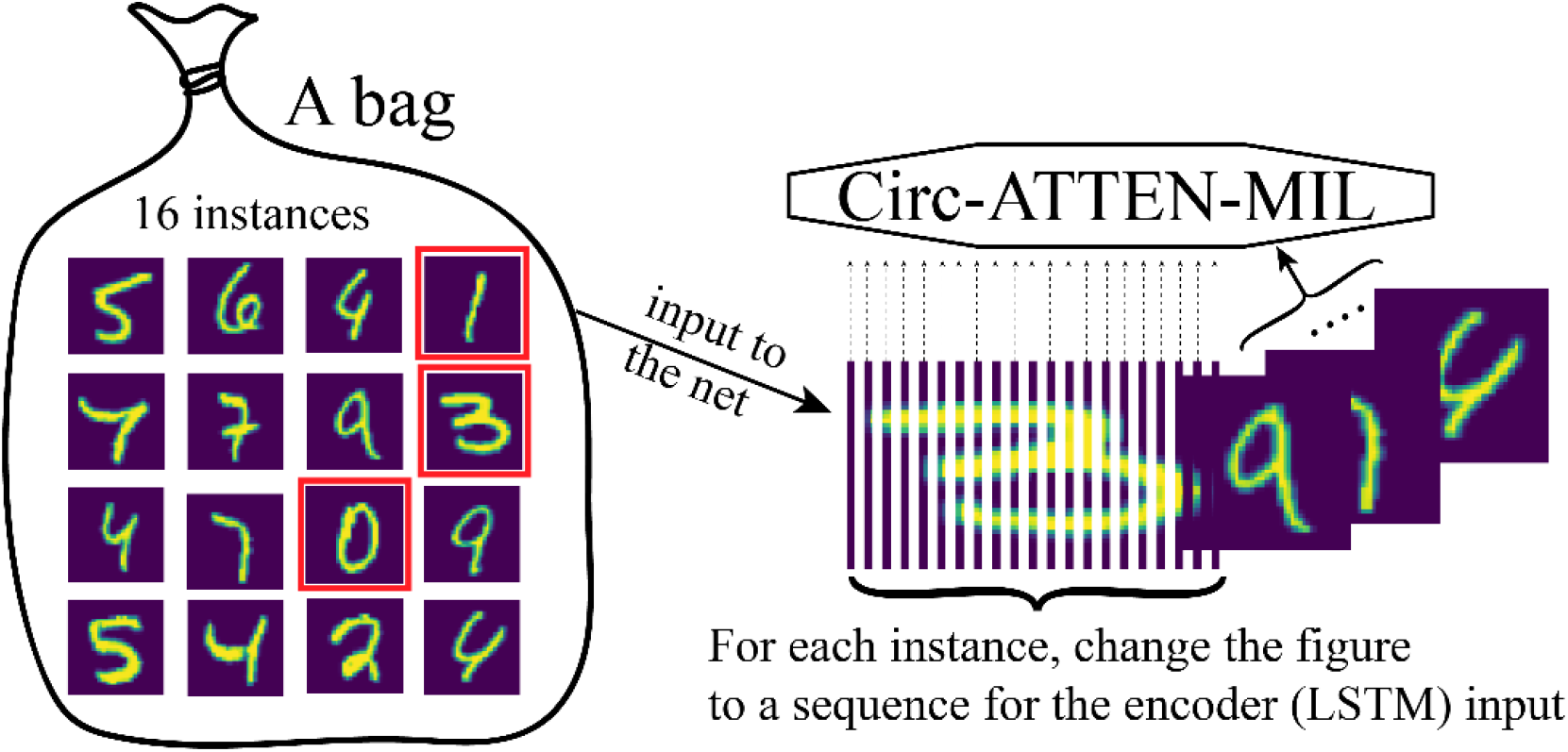
Handling of handwritten numbers dataset for feeding into Circ-ATTEN-MIL.

### Fusion model

The ‘weighted feature’ (the penultimate layer) of Circ-ATTEN-MIL was extracted as the sequence feature defined by the model. The other features were calculated using the extraction methods of RCM features and conservation features in CircDeep. Combining these three types of features (sequence feature: 100; RCM feature: 40; conservation feature: 23), a four-layer MLP (Multi-Layer Perceptron) network (163-80-20-1 (the output layer is a sigmoid-activated neuron)) was constructed as a fusion model.

### Evaluation criteria

We evaluated the model performance by classification accuracy, sensitivity, specificity, MCC (Matthews correlation coefficient), and F1 score (formulated as follows).

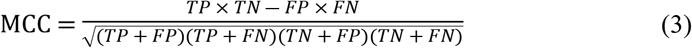

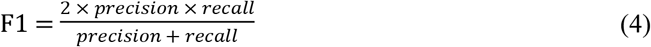

### Extraction of highly attention sequence splices

As the attention score was applied to the encoder features of each instance, we assigned the same scores to the sequence of the instance, and collapsed the weighted sequences according to the inverse of the slider rule (Fig.4), and extracted the sequence fragment (with certain length: >7) with the higher attention score (after scaling to between 0 and 1: >0.6), which served as the high attention sequence splices.

**Figure 4.**
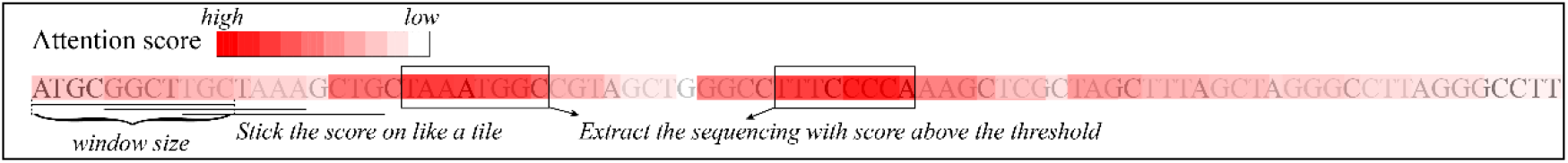
Illustrations of extraction of highly attention sequence splices

### Motif enrichment

MEME software^[28]^ was utilized to perform motif enrichment tasks. In MEME environment, classic mode was selected to enrich motifs in RNA sequences between 6 and 50 lengths (The code is: *meme RNA.fasta -rna -nostatus -mod zoops -minw 6 -maxw 50 -objfun classic -markov_order 0*).

### Result 1: Dataset description

The sequence length distribution and base proportion between circRNAs and other lncRNAs (In training set) were very similar (Fig.5), which illustrated that the simple features between the two-type sequences were comparable and the model feeding with raw sequences was hard to accomplish the identification task by these simple features.

**Figure 5.**
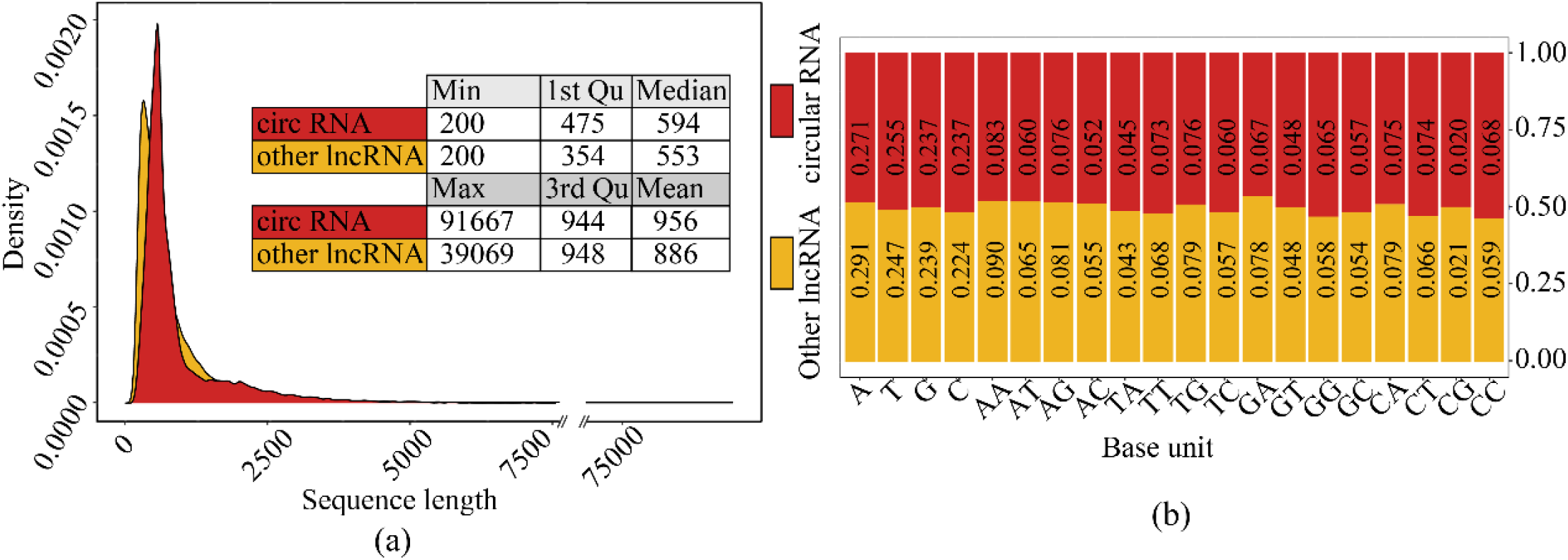
The comparison of simple features of sequences between the two-type sequence set: (a). Sequence length distribution comparison. (b). sequence composition comparison.

### Result2: Model architecture

In instance extraction, the window size was set to 70, sliding step was set to 5 (Fig.1). In the encoder block, it consists of one embeding_15_4 layer and two bi-direction LSTM_4_150 layers. The final step outputs of both directions were concatenated, and via an FCN_300_100 layer, the instance feature (C_100) was obtained. In the attention block, the C_100 features of each instance were accepted as key values. After an FCN_100_30, an FCN_30_1 layer, the dimension for each instance was reduced to 1 (attention value). A softmax layer was utilized to normalize the attention value for each instance, and then the normalized attention score was yield. Finally, the classifier block accepting all instances’ weighted C_100 feature, through a fully connected layer and a sigmoid neuron, outputted the identification probabilities (Fig.2).

### Result3: Model training and identification evaluation

We used the binary cross-entropy loss function to calculated loss and trained the models with the adam optimization algorithm (learning rate is 0.0002; betas = (0.9, 0.999); weight decay is 10e-5). Balancing the accuracy and over-fitness, we chose the model trained at the 70th epoch as the final model and plotted the ROC curve (Fig.6). As a result, the performance of the model training had strong identification efficiency (Train AUC=0.99; Validation AUC=0.97; Test AUC=0.97). Subsequently, multiple evaluation criteria were employed to test the model (Table 1), and these metrics also validated that the model has a high degree of robustness.

**Figure 6.**
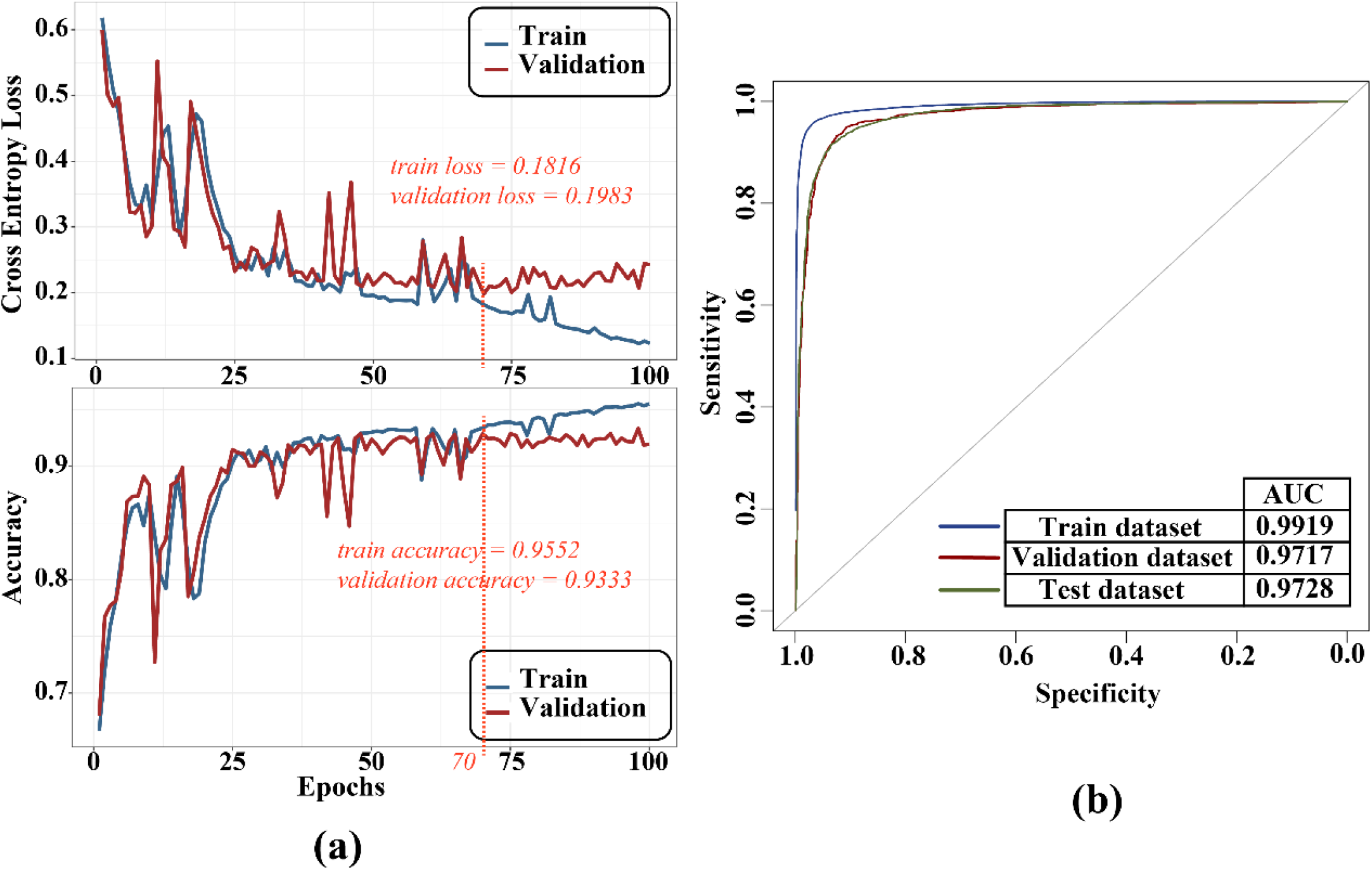
Training process (a) and ROC curve (b).

**Table 1.** The evaluation for classification task

|  | *Accuracy* | *Sensitivity* | *Specificity* | *Precision* | *MCC* | *F1* |
| --- | --- | --- | --- | --- | --- | --- |
| ***Train*** | 0.9552 | 0.9662 | 0.9547 | 0.9713 | 0.9194 | 0.9687 |
| ***Validation*** | 0.9333 | 0.9485 | 0.9092 | 0.9433 | 0.8291 | 0.9459 |
| ***Test*** | 0.9284 | 0.9396 | 0.9039 | 0.9393 | 0.8435 | 0.9394 |

### Result4: Comparison with other algorithms

This model was compared with ACNN-BLSTM in CircDeep^[13]^, which took the sequence as input without the feature from the secondary structure and conservation score of the sequence. In Circ-ATTEN-MIL, the input was full-length raw sequences. While in ACNN-BLSTM, the input was the padded triplet sequences (the base triplet was transformed to a 40-dimension vector by word2vec; the input length was padded to 8000). The comparison results showed that our final model was better under the three metrics (Table.2). Finally, we incorporated the RCM and conservation features which used in CircDeep model to build a fusion model (Methods), and successfully improved the discriminative power of the final model.

**Table 2.**
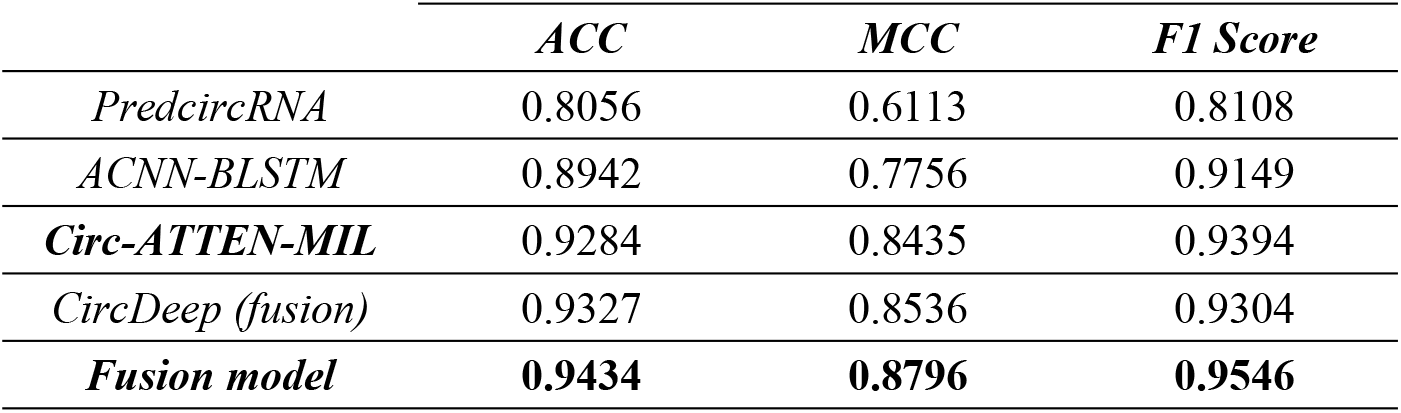
The comparison results

### Result5: Attention score employed for identifying determining factor

To verifying the representational power of the attention score, we used the handwritten numbers dataset to visualize the known determining factor with the produced attention score. Two model (In encoder block: 2 LSTM_28_10, FCN_10_10; in MIL block: FCN_10_5, FCN_5_1) was trained in this part, one (model 1) with 0 and another (model 2) with 0, 1, 3 as determining number (a bag contains determining number instances was treated as positive sample). The training was stopped after the accuracy exceed 0.90 (around 10 epochs). We visualized the attention score with the matched instances and discovered that the attention score identifies well whether the bag contains a single determinant, multiple identical determinants, or multiple different determinants (Fig.7). Statistics on determining numbers identification showed a very low percentage of false identifications, and although there was a certain unrecognized rate, the identified numbers had a very high confidence level (>99%).

**Figure 7.**
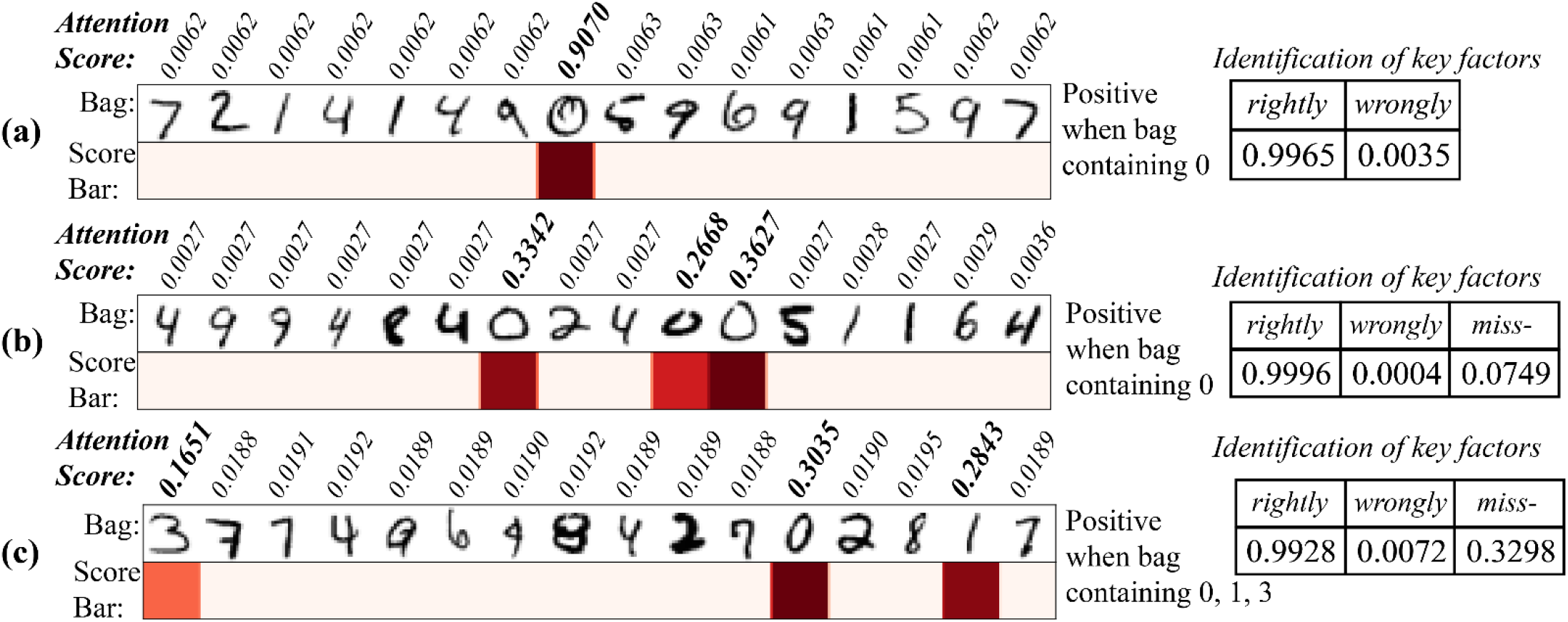
Attention score for identifying the determining numbers. (a). single determinant (model 1); (b). multiple identical determinants (model 1); (c). multiple different determinants (model 2). (left panel: attention score bar; right panel: the rightly and wrongly identify events and miss-identify events).

### Result6: Motif enrichment from high attention sequence

The high attention sequences were extracted for all correct identification circRNAs transcripts. Most of the high attention sequences were between 8-40 in length, and the count of the attention sequences for each transcript was around 4 (Fig.8), which validates our initial assumption that the meaningful features were sparse. All high attention sequences were used for motif enrichment, and multiple validated motifs were yield (Table 3).

**Figure 8.**
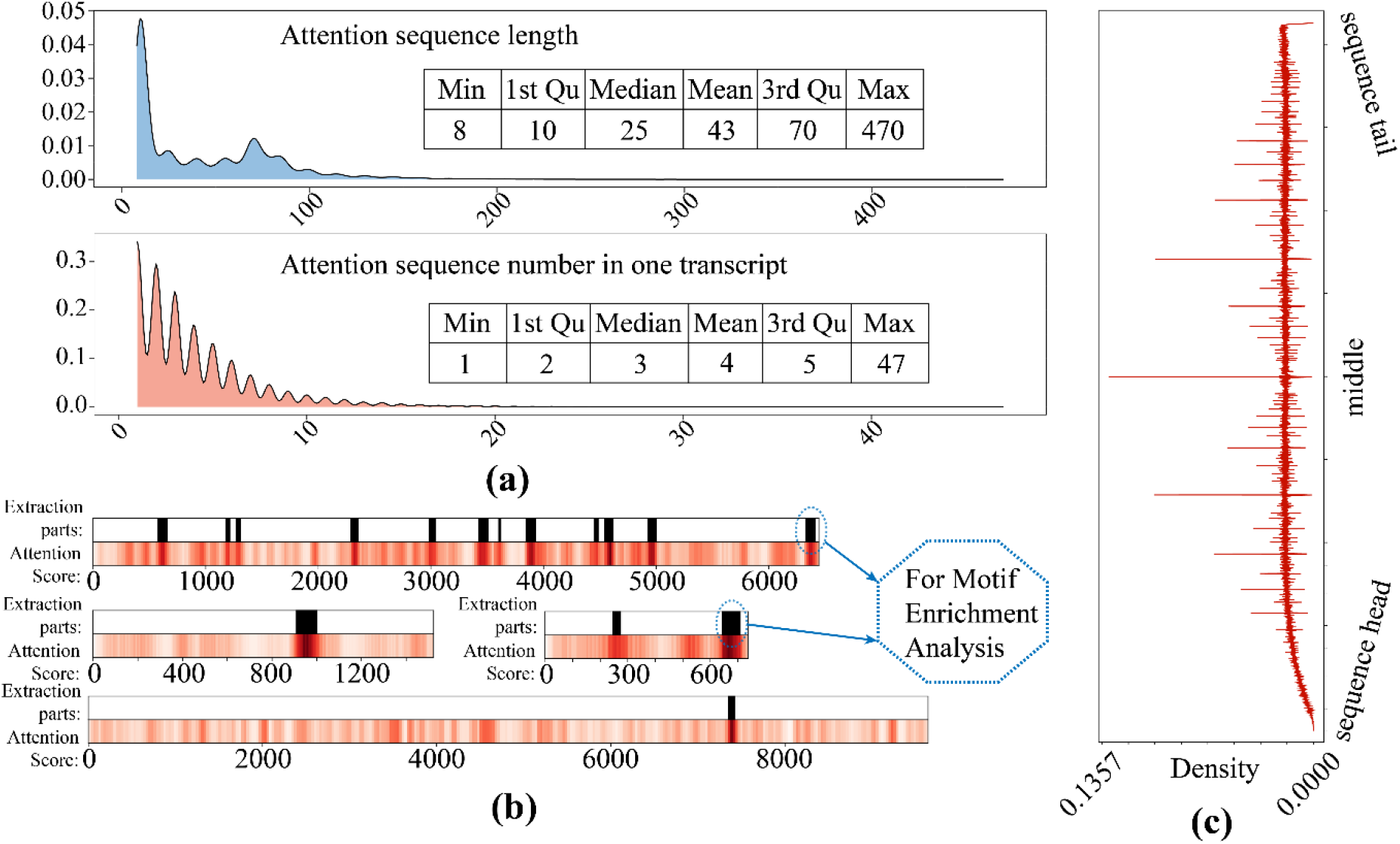
The high attention sequence distribution. (a) the length distribution (upper) and the attention sequence number for each transcript distribution (lower); (b). the extraction of attention sequence for motif enrichment; (c). density distribution of attention loci on all sequences.

**Table 3.**
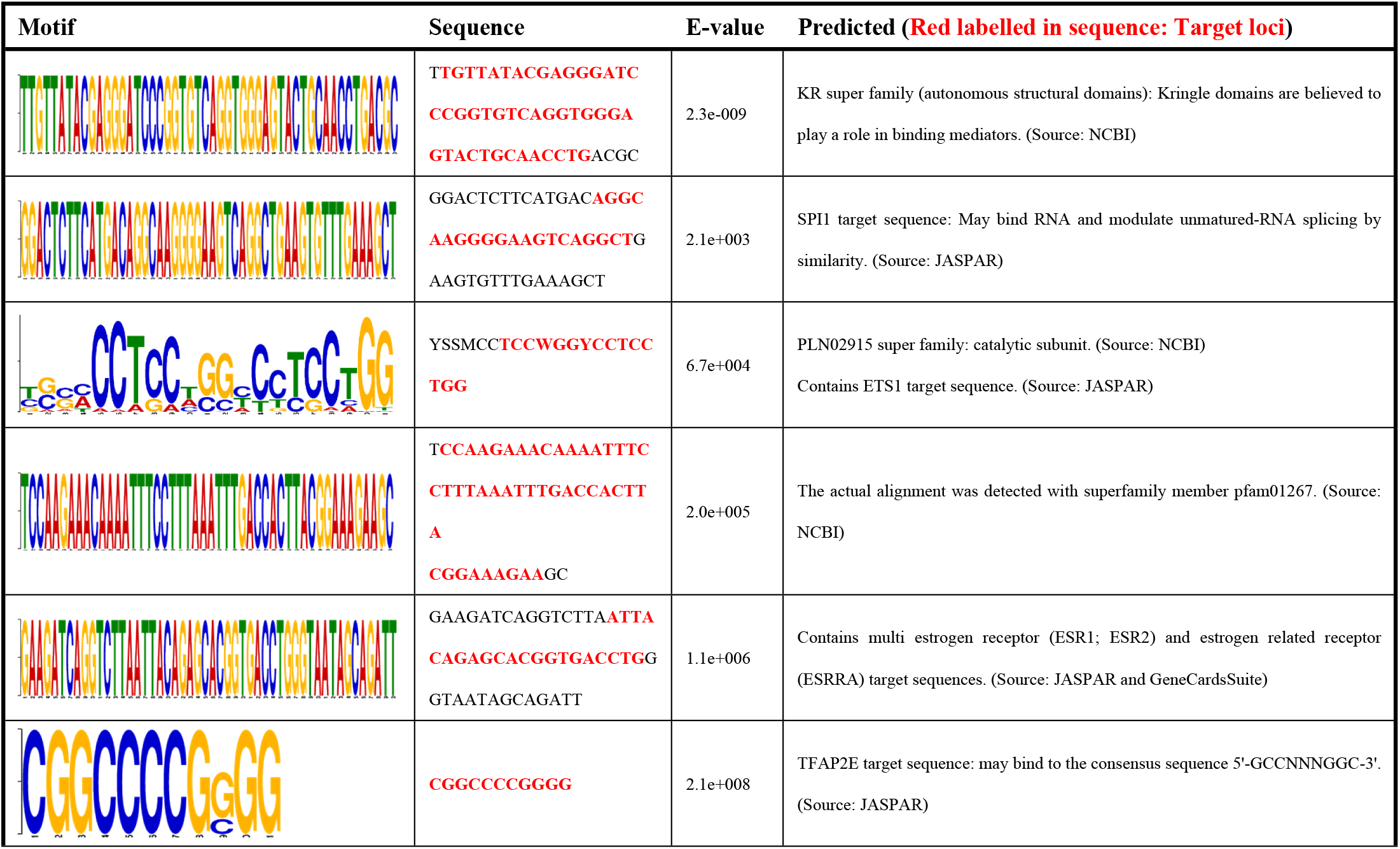
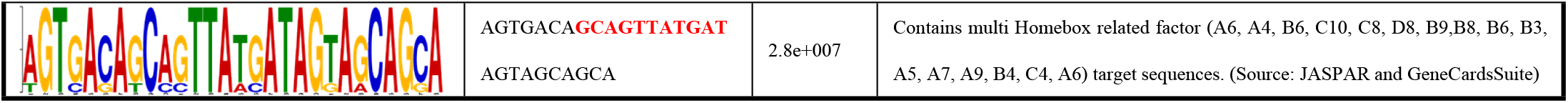
motif enriched from the sequence

## Discussion

In this project, we designed a deep learning network architecture suitable for learning gene sequence features and implemented the model to accomplish the circRNA identification task. And based on the attention score produced by the model, a large number of key sequence loci for circRNA recognition were extracted. Following the motif enrichment analysis, some possible key motifs for circRNA formation were identified.

The post-transcriptional modifications and a variety of related functions of transcripts are encoded in their sequence^[29]^. Thus, a sequence contains a large number of key loci responsible for each of the processes^[30]^. For machine learning models, which often responsible for discriminating a single function, such as loop formation, the entire sequence can be too redundant and the meaningful features are too sparse. From another viewpoint, the learning-by-sequence task is similar to multiple instance learning (MIL)^[20, 27, 31]^, that is, for weak label learning problems with sparse features. We changed the convolutional blocks commonly used in the MIL model for feature extraction to an RNN block that is more suitable for sequence learning^[32]^, and used the attention mechanism^[21, 33]^, which has stronger representation capability, as the MIL layer. The results demonstrate the validity of the structure and the great potential value of the attention mechanism.

For this circRNA identification task, data were collected from the validated reference sequence with high confidence^[22, 23]^. While there are certain problems that the sampling rate was too low. If a single gene is assumed to be a single distribution (which may actually be a set of genes), and the use of a reference sequence causes only one sample to be collected for a single distribution, the sampling rate can be considered to be relatively low. Therefore, if multiple actual sequences can be collected for a single gene, which implies that there may be a variety of mutations in non-relevant features of multiple sequences, while relevant features are more conservative, the increased sampling rate must enhance the model’s learning of the features and improve its discriminative power. Considering that data collection is more difficult^[34]^, it is worthwhile to explore to improve the effectiveness of the model by trying some data augmentation methods.

The instance is extracted by a moving slider, which can only extract the continuous regional features in the sequence. However, sequences form higher-level stereo structures in space^[35]^, so the key feature can be the combinations of sequences that are far apart. Considering this possibility, adding more mechanisms for instances extraction and combination, to make a single instance can contain multiple combinations of distant sequences, may further improve the discriminative effectiveness as well as the potential representational value of this network structure.

The model can be used for more than just the identification of circRNAs. Since only the original sequence is required as input, the network structure can be used for learning other sequence-related tasks by simply changing the resultant events.

Because of its representation capability, it can be used to discover key sequences for different tasks and provide a basis for other relevant research.

## Conclusions

Circ-ATTEN-MIL was designed and used for circRNA identification, and it outperformed other deep learning models currently used. The model utilized the MIL-attention network architecture, which took the complete RNA sequence as input and not only carried out the discriminant probability of circRNA identification, but also outputted the score of the importance of each instance, which could be used for identifying the critical part of a sequence for model judgment and would be able to provide some insights for basic research in related fields.

## Declarations

### Ethics approval and consent to participate

Not applicable (No human participation).

### Consent for publication

All authors agree to publication.

### Availability of data and material

The data and code are available in https://github.com/liuyunho/Circ-ATTEN-MIL, and any other requirement can contact the corresponding author.

### Authors’ contributions

Conceptualization: Y.L., G.L. and Q.F.; methodology: Y.L., Q.F; network design: Y.L.; validation: Y.L., X.P. and C.Z.; writing—original draft preparation: Y.L. and X.P.; visualization: Y.L. and X.P.; funding acquisition: L.L.

